# Shade-avoidance responses become more aggressive in warm environments

**DOI:** 10.1101/710004

**Authors:** Sofía Romero-Montepaone, Sofía Poodts, Patrick Fischbach, Romina Sellaro, Matias D. Zurbriggen, Jorge J. Casal

## Abstract

When exposed to neighbour signals, competitive plants increase the growth of the stem to reduce the degree of current or future shade. Plants can experience similar neighbour cues under different weather conditions and the aim of this work is to investigate the impact of daily average temperature and irradiance and thermal amplitude on the magnitude of shade-avoidance responses in *Arabidopsis thaliana*. For this purpose, we first generated a growth database and elaborated under controlled conditions a model that predicts hypocotyl growth during the photoperiod as a function of the light-modulated activity of the main photo-sensory receptors (phytochromes A and B, cryptochromes 1 and 2), temperature-modulated activity of phytochrome B and an independent temperature input. Thermal amplitude (lower temperatures during the morning and afternoon that at midday) reduced growth of genotypes with normally fast morning growth, and this effect was incorporated to the model. Thermal amplitude also decreased the abundance of the growth-promoting transcription factor PHYTOCHROME-INTERACTING FACTOR 4. The model predicted growth in the field through different seasons with reasonable accuracy. Then, we used the model in combination with a worldwide dataset of current and future whether conditions. The analysis predicted enhanced shade-avoidance responses as a result of higher temperatures due to the geographical location or global warming. Irradiance and thermal amplitude had no effects. These trends were also observed for our local growth measurements. We conclude that, if water and nutrients do not become limiting, warm environments enhance the shade avoidance response.

## 1 INTRODUCTION

Stem growth can affect the yield of agricultural crops (Foulkes *et al.* 2011). Excessive growth enhances the vulnerability to wind and the associated risk of lodging, as well as the competition for resources between the stem and harvestable organs. Variations in these factors were of fundamental importance for the success caused by the introduction of dwarfing genes during the breeding process that generated the green revolution. At the other end of the scale, short plants can also be sub-optimal as the resulting canopy architecture involves strong overlap among leaves and poor penetration of the light. This condition may result in upper leaves with photosynthesis saturated by light simultaneously with lower strata with negative net carbon dioxide exchange.

The light environment has a major influence on stem growth. Due to the optical properties of the leaves, which absorb strongly in the UV, blue and red wavelengths while transmitting and reflecting more in the far-red range, the presence of nearby vegetation reduces the red / far-red ratio even if the plants are close-by without shading each other, and reduce the irradiance as mutual shading becomes more intense (Franklin 2008; Martínez-García *et al.* 2010; Casal 2013; Ballaré & Pierik 2017). These neighbour signals reduce the activity of photo-sensory receptors that control stem growth. The low red / far-red ratios and/or the reduced red irradiances of canopy shade are perceived by phytochrome B (phyB) in *Arabidopsis thaliana* (Robson, Whitelam & Smith 1993; Sellaro *et al.* 2010; Trupkin, Legris, Buchovsky, Rivero & Casal 2014), *Brassica rapa* (Devlin, Rood, Somers, Quail & Whitelam 1992) cucumber (Ballaré, Casal & Kendrick 1991; López Juez *et al.* 1992) and tomato (Casal 2013). The low blue-light irradiances and low blue / green ratios caused by neighbours are perceived by cryptochrome 1 (cry1) and cry2 (Sellaro *et al.* 2010; Keller *et al.* 2011). Increasing intensities of neighbour cues initiate the so-called shade-avoidance responses, including enhanced stem growth, mainly by reducing the activities of phyB and cry1. At high irradiances (as those observed in the field), phyA becomes an effective sensor of red irradiance in addition to far-red light and therefore, phyA also contributes to the inhibition of stem growth by progressive reductions in canopy shade (Franklin, Allen & Whitelam 2007; Sellaro *et al.* 2010). Finally, the UV-B RESISTANT 8 (UVR8) receptor perceives brief interruptions of canopy shade to reduce stem growth (Moriconi *et al.* 2018). Major points of signalling convergence downstream these photo-sensory receptors include PHYTOCHROME INTERACTING FACTORS (Leivar & Quail 2011; Hornitschek *et al.* 2012) and CONSTITUTIVE PHOTOMORPHOGENIC 1 (Lau & Deng 2012; Pacín, Semmoloni, Legris, Finlayson & Casal 2016).

In addition to its role in the perception of the light environment, phyB is also able to sense temperature (Jung *et al.* 2016; Legris *et al.* 2016). While high, compared to low, red / far-red ratios increase the proportion of phyB in its active conformation warm temperatures have the opposite effect via thermal reversion from the active to the inactive phyB conformer. When irradiance is not high, phyB is in a steady state that depends strongly on temperature (Sellaro, Smith, Legris, Fleck & Casal 2019). The blue-light receptor phototropin (Fujii *et al.* 2017) has also been shown to sense temperature but this photo-receptor only has a transient effect on straight stem growth (Folta & Spalding 2001).

Plants can eventually experience similar neighbour cues under different weather conditions. The overall irradiance can be affected by solar angle and clouds, and air temperature patterns are very dynamic. The aim of this work was to elucidate the degree of impact of weather conditions (specifically irradiance patterns, mean temperature and thermal amplitude) on the response of stem growth to plant neighbour cues. This issue is particularly important in the scenario of global warming and climate change because crops where stem growth is adjusted to current weather conditions could become sub-optimal in terms of architecture in the future. A reductionist approach could simply involve testing the effect of similar neighbour signals in combination with different temperatures and overall irradiances under controlled environments. The latter type of experiments has already been conducted in the past (Wall & Johnson 1982; Mazzella, Bertero & Casal 2000; Weinig 2000; Halliday & Whitelam 2003; Patel *et al.* 2013). Although informative, such approach bears serious limitations to predict plant responses under natural conditions for two reasons. One is that the natural environment is more complex than the conditions typically used in growth cambers. For instance, light and temperature fluctuate diurnally and this pattern is difficult to simulate in combination with many light conditions. Second, some environmental variables such as overall irradiance and temperature do not fluctuate fully independently in nature and the use of simulated combinations do not necessarily reflect those that a plant is more likely to find in the field. Therefore, to address this issue we followed a more sophisticated approach, where we conducted experiments under controlled conditions to extend a stem growth model based on phyB activity (Legris *et al.* 2016) to incorporate the action of cry1, cry2 and phyA. We also analysed the need to incorporate ad-hoc model terms to account for the effects of diurnal thermal and irradiance fluctuations. Then, we generated a growth database by cultivating *A. thaliana* plants in the field under sunlight and two levels of shade, and tested against this data the predictions of the model parameterised under controlled conditions. Since the model predicted stem growth with reasonable accuracy, we used it in combination with weather data from different regions of the Earth and models that predict future trends to evaluate the impact of realistic weather conditions on the shade avoidance response.

## 2 MATERIALS AND METHODS

### 2.1 Plant material

We used seedlings of *Arabidopsis thaliana* of the wild type Columbia background in all the experiments. Wild type (WT) plants and the *phyA-211, phyB-9, phyA-211 phyB-9 cry1-304, cry2-1* and *cry1-304 cry2-1* mutants (for references see Sellaro, Pacín & Casal 2012) were included in all the experiments, except those aimed to model the action of either phyA or cry1 and cry2, which included only the relevant mutants. For all the experiments, 10-15 seeds per genotype were sown on 8 ml of 0.8 % agar in clear plastic boxes and kept in the dark at 4°C for 4 days. Then, they were transferred to white light conditions for 3 d in a growth room at 22°C with a photoperiod of 10 under 100 µmol. m^-2^. s^-1^ (400-700 nm) provided by a mixture of fluorescent and halogen lamps (red / far-red ratio = 1.1). At the time of initiation of the photoperiod of the fourth day, the seedlings were transferred to the corresponding experimental light conditions.

### 2.2 Hypocotyl length increment

For field experiments and all indoor experiments under constant conditions pictures of the seedlings were taken at the beginning and at the end of the photoperiod of the fourth day. For experiments with fluctuating temperatures, pictures were taken every 2.5 during the photoperiod of the fourth day. For experiments with fluctuating light, pictures were taken at the beginning, at the middle and at the end of the photoperiod of the fourth day. Hypocotyls were measured using image processing software (Sellaro, Hoecker, Yanovsky, Chory & Casal 2009) and length increments were divided by the duration of the period to obtain growth rates. Data obtained in experiments under controlled conditions are shown in Table S1 and those obtained in the field are shown in Table S2. When using growth data to either adjust or test the growth model, the hourly growth rates were converted to length increments during a period of 9 h (by multiplying by 9) for consistency with the original model (Legris *et al.* 2016).

### 2.3 Light treatments under controlled conditions

To model the contribution of phyA (see below Fig. 1a-b) we used five different mixtures of red and far-red light provided by 150 W incandescent R95 lamps (Philips) in combination with a water filter, a yellow, orange and red acetate filter set (LEE filters 101, 105 and 106, respectively), either alone or plus a green acetate filter (LEE filters 138, 121 or 089) or six blue acrylic filters (Paolini 2031, Buenos Aires, Argentina). Scans of the light fields were obtained with a spectroradiometer (USB400, Ocean Optics, USA) The red / far-red ratios were: 1.09, 0.44, 0.20, 0.07 and 0.001 and the irradiances ranged from 63 to 834 μmol. m^-2^. s^-1^ (spectra are shown in Fig. S1a). Spectral data were used in combination with photoconversion cross section data (Mancinelli 1994) to obtain the rates of photoconversion from Pr to Pfr (k1) and from Pfr to Pr (k2). To model the contribution of cry1 and cry2 (see below Fig1c-d) blue light was provided by light-emitting diodes (the spectrum is shown in Fig. S1b).

**Figure 1.**
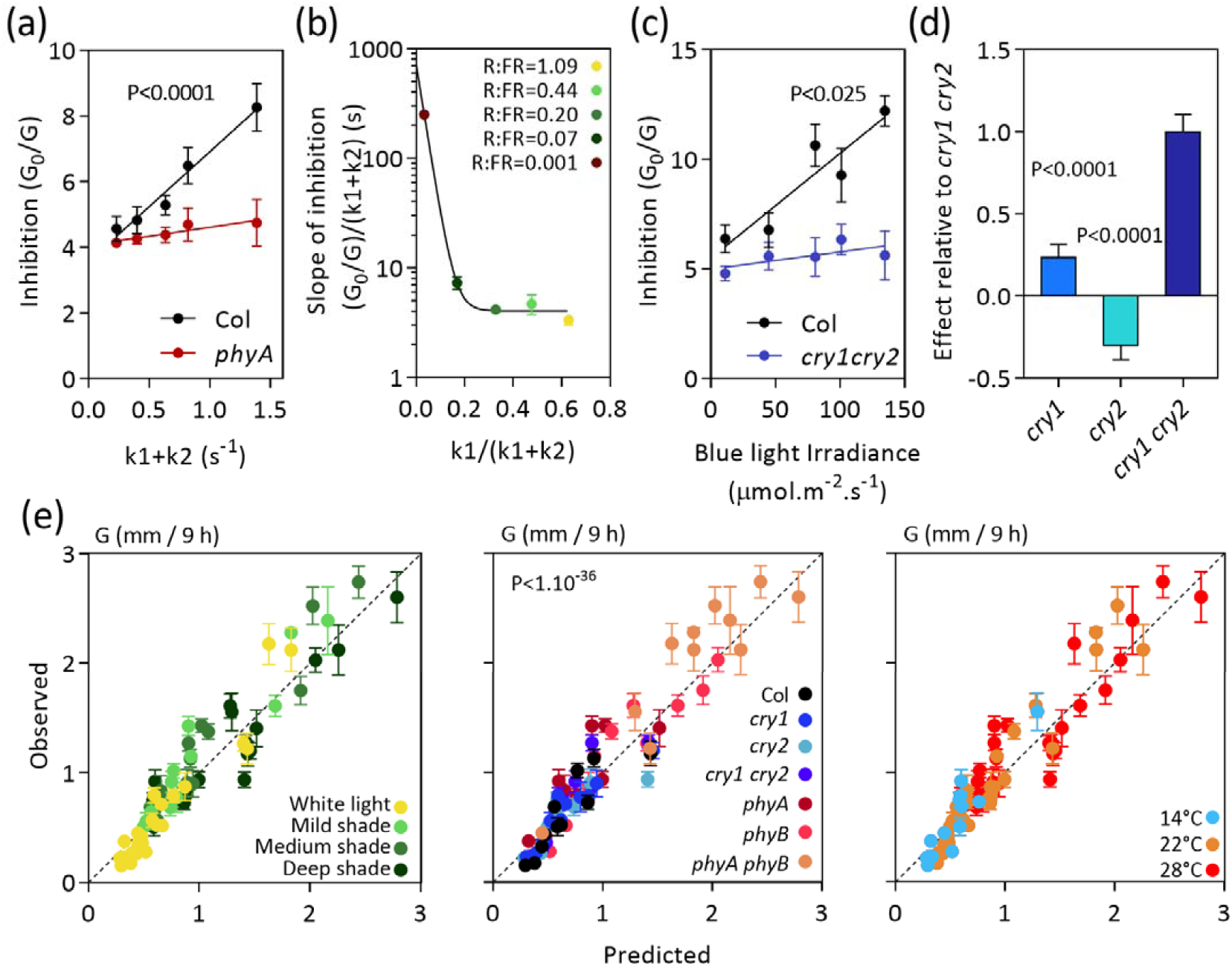
Modelling the contribution of phyA, cry1 and cry2 to the control of hypocotyl growth. (a) Growth inhibition of the WT and the *phyA* mutant in response to the irradiance of a red plus farred light mixture (red / far-red ratio= 1.09), represented by the sum of photo-conversion rates (k_1_+k_2_). Data are means and SE of five replicate boxes of seedlings. (b) Relationship between the difference in slope of the response to k_1_+k_2_ in the WT and the *phyA* mutant and the k_1_/(k_1_+k_2_), which increases with the proportion of red light in the of the red plus far-red light mixture. (c) Growth inhibition of the WT and the *cry1 cry2* double mutant in response to the irradiance of blue light. Data are means and SE of four replicate boxes of seedlings. (d) Relative contribution of cry1 and cry2. Growth of *cry1* or *cry2* mutants relative to the growth of the *cry1 cry2* mutant. (e) Observed values of hypocotyl growth (G, Table S1) in seedlings of seven genotypes grown under different combinations of light/shade and temperature versus the growth values predicted by the model where the term based on irradiance (Legris *et al.* 2016) is replaced by the specific contributions of phyA, cry1 and cry2. The different light conditions, genotypes and temperatures are color-coded to show that the relationship between observed and predicted values is not biased for any of these factors (within the range tested here).

To test the model under stable conditions of light and temperature, simulated sunlight was provided by a combination of fluorescent and halogen lamps (100 μmol. m^-2^. s^-1^, between 400 and 700 nm) and increasing degrees of shade were simulated by the combination with different green acetate filters (LEE filters 138, 121 or 089 for mild, medium and deep shade, respectively). The red / far-red ratios were: 0.76, 0.41 and 0.12 and the irradiances between 400 and 700 nm ranged from 18 to 70 μmol. m. s (spectra are shown in Fig. S1e). Different growth chambers were used for the different temperature conditions (14°C, 22°C and 28°C).

For experiments with temperature fluctuations (see below Fig. 2) we used a growth chamber (Percival E-308) with fluorescent light tubes (30 μmol. m^-2^. s^-1^ between 400 and 700 nm, the spectrum is shown in Fig. S1c) and simulated real temperature patterns obtained from climate stations at three different locations in Argentina (*25 de Mayo, Ingeniero Jacobacci and Puerto Pirámides*) for a date when photoperiod was 10 h. The three cases show similar medium temperature but different thermal amplitudes.

**Figure 2.**
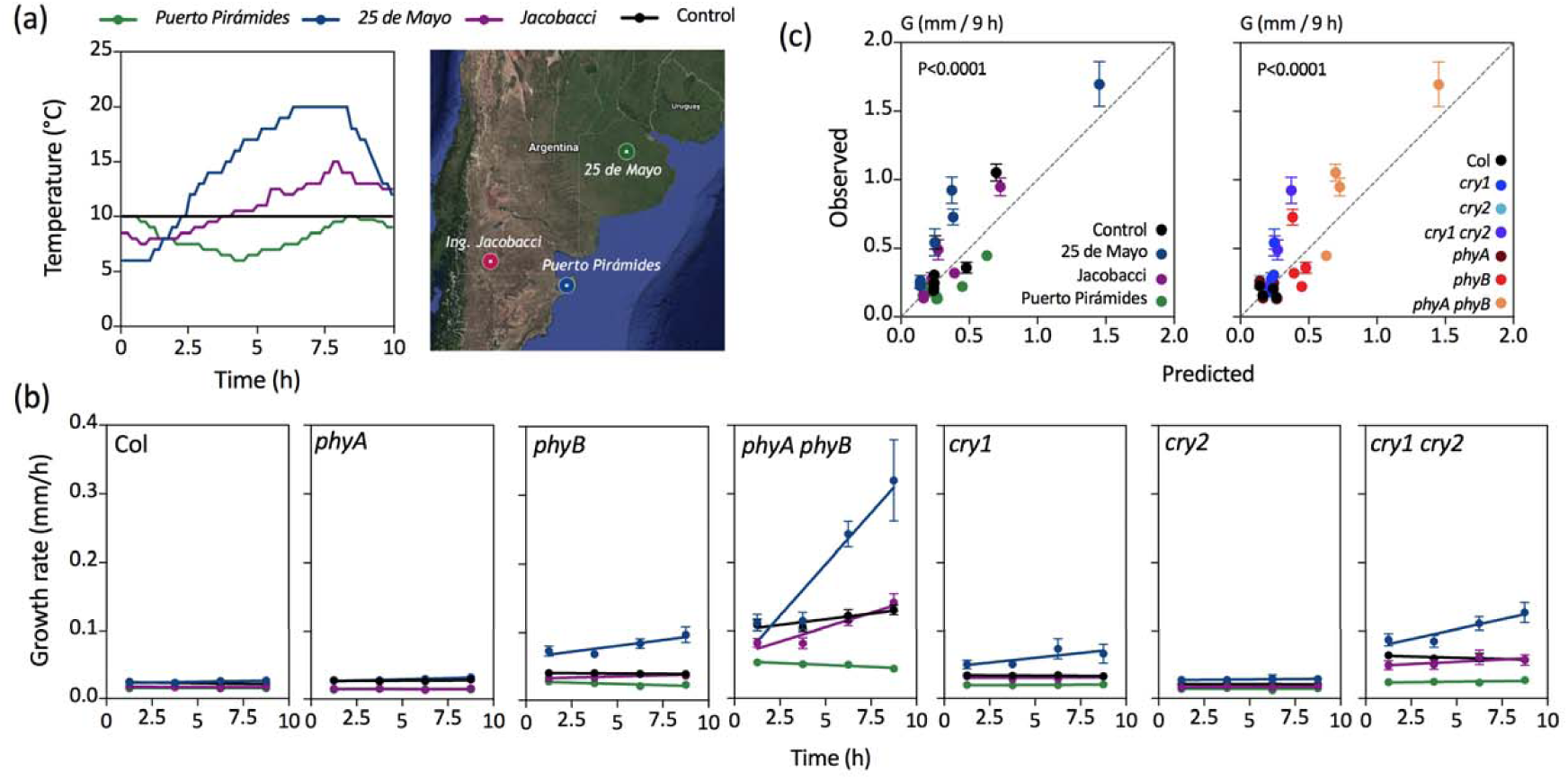
Thermal amplitude reduces stem growth. (a) Temperature patterns used in the experiments. (b) Growth rates under the different temperature regimes (Table S1). Data are means and SE of four replicate boxes of seedlings. (c) Observed values of hypocotyl growth (G) versus the growth values predicted by the model incorporating a term for thermal amplitude (Table S3h). The different temperature patterns and genotypes are color-coded to show that the relationship between observed and predicted values is not biased for any of these factors (within the range tested here).

For experiments with light fluctuations (see below Fig. 3), we used a LED panel with 3 light channels: red, far-red and blue (spectra are shown in Fig. S1d). For the control, each one of the three channels was kept constant at 9 μmol. m^-2^. s^-1^. For the blue light treatment the red and far-red channels were kept constant at 9 μmol. m^-2^. s^-1^ and blue light intensity was increased 20% every hour (starting with 3.6 μmol. m^-2^. s^-1^) during the first half of the photoperiod (when it reached maximum intensity, 18 μmol. m^-2^. s^-1^) and then decreased symmetrically, i.e. 20% every hour during the second half of the photoperiod. The same pattern and intensity values were used for red plus far-red treatment, where the blue channel was kept constant and the red and far-red channels were shifted during the photoperiod. For the blue plus red plus far-red treatment all the channels were shifted during the photoperiod following the same pattern.

**Figure 3.**
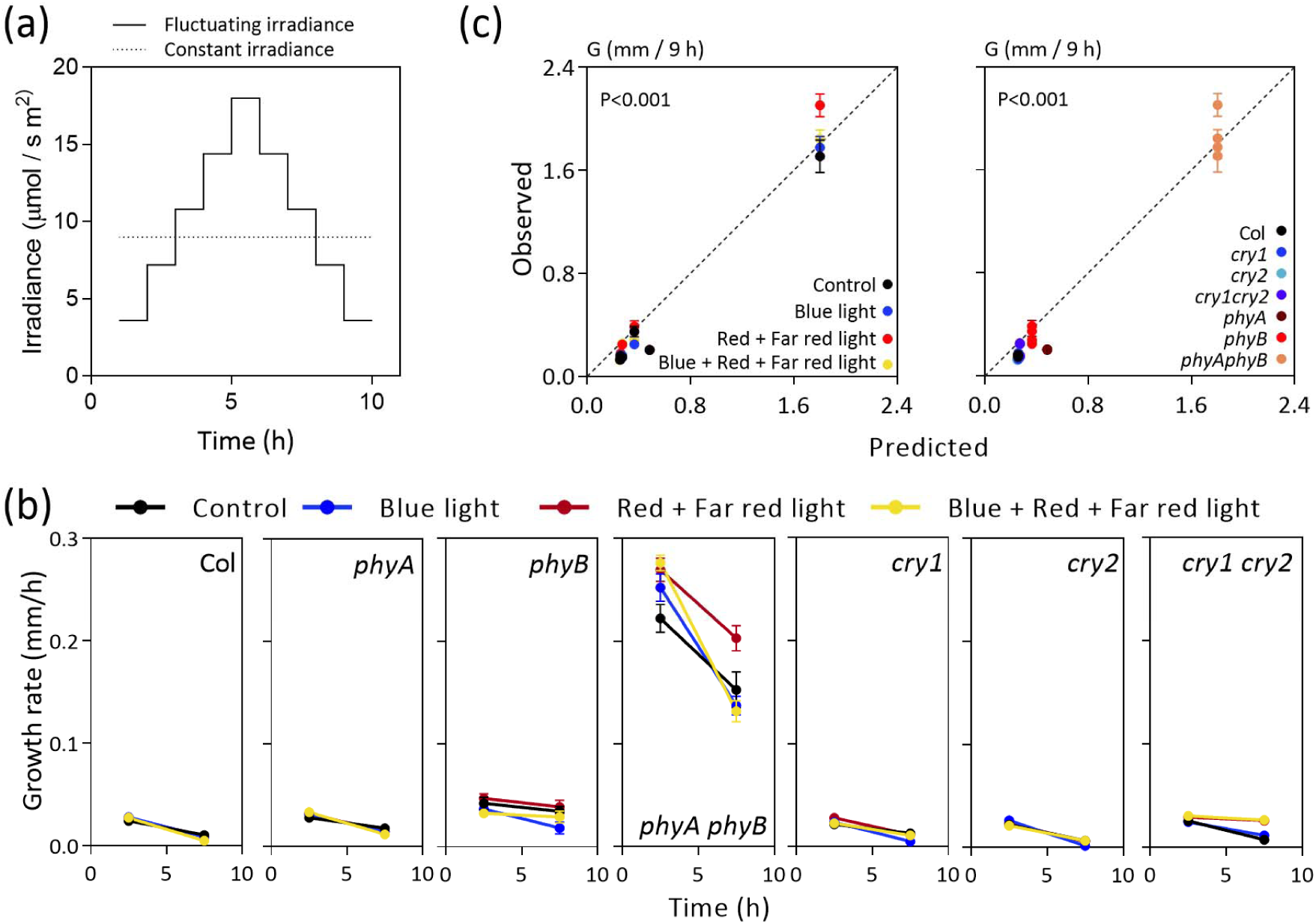
Diurnal fluctuations in irradiance have minor effects on stem growth. *(a)* Light patterns used in the experiments. (b) Growth rates under the different light regimes (Table S1). Data are means and SE of three replicate boxes of seedlings. (c) Observed values of hypocotyl growth (G) versus the growth values predicted by the model (Table S3h). The different light patterns and genotypes are color-coded to show that the relationship between observed and predicted values is not biased for any of these factors (within the range tested here).

### 2.4 Field experiments

On the fourth day light grown seedlings of all the genotypes were transferred either to unfiltered sunlight or under one of two different canopy shade conditions (mild and deep shade) in the experimental field of *Facultad de Agronomía*, University of Buenos Aires (latitude -34.5914, longitude -58.4798). Ten different experiments were performed at different dates, starting in June (winter) and ending in February (summer) to obtain arranges of different light and temperature combinations. Temperature inside the growth boxes that contained the seedlings was recorded hourly by using a data-logger (DS1922L-F5# Thermochron iButton). Photosynthetically-active radiation, blue, red and far-red light were recorded hourly by using a 4-channel light sensor (SKR 1850, Skye Instruments).

### 2.5 Weather data

To investigate the impact of diurnal temperature fluctuations on hypocotyl growth, we searched a local weather database (*SIGA, Sistema de Información y Gestión Meteorológica INTA, Argentina,* http://siga2.inta.gov.ar/). We identified locations and dates where the photoperiod was 10 h. Then, we selected tree cases where mean temperature was similar (close to 10°C) and thermal amplitude was contrasting. One of the locations (*25 de Mayo*) showed considerable thermal amplitude, the second (*Ingeniero Jacobacci*) showed small thermal amplitude, and the third (*Puerto Pirámides*) showed thermal inversion caused by the important oceanic influence.

To apply the growth model at a global scale of current weather, maximum temperature, minimum temperature and solar radiation data for 100 different locations were obtained from the CCAFS-Climate data portal (http://www.ccafs-climate.org). The processing of datasets was conducted by using GIS software (QGIS 3, http://www.qgis.org/). The data correspond to dates when the photoperiod is close to 10 h (this excludes latitudes where days are longer throughout the year).

For future projections (year 2080) of weather conditions, the climate model mpi-echam5 (http://ccafs-climate.org/data_spatial_downscaling/) and the emission scenario A1B were used. The latter scenario predicts an integrated world characterised by rapid economic growth, a population that reaches 9 billion by 2050 and then declines gradually, and the rapid development of alternative energy sources that facilitate increased economic growth while limiting and eventually reducing carbon emissions. The starting point (current weather conditions) was defined by the conditions obtained from the CCAFS-Climate data portal, as described above.

### 2.6 Bioluminiscence assay

For bioluminescence assays, we exposed 3-day-old light-grown seedling of the Columbia wild-type carrying the *PIF4:PIF4-LUC* transgene to a photoperiod with irradiance or temperature manually adjusted hourly to obtain a peak at midday, compared to a control under constant light and temperature. Seeds carrying the bioluminescent PIF4 reporter were kindly provided by Salomé Prat (Centro Nacional de Biotecnología, Madrid, Spain) and will be described in further detail elsewhere. In the morning, light increased linearly from 41 μmol. m^-2^. s^-1^ during the first hour to 230 μmol. m^-2^. s^-1^ during the fifth hour. Similarly, temperature increased linearly from 13 °C during the first hour to 27 °C during the fifth hour. In both cases, the opposite trend was applied during the afternoon. The control was exposed to 135 μmol. m^-2^. s^-1^ and 20 °C throughout the 10 h photoperiod. Twenty-four hours before starting luciferase readings, 20 µL of 0.5 mM D-luciferin were added to each well. Luciferase activity was detected with a Centro LB 960 (Berthold) luminometer.

## 3 RESULTS

### 3.1 Basic growth model under stable light and temperature

The model developed by Legris et al. (2016) predicts the average hypocotyl growth of a 3-day-old light-grown seedling during the photoperiod as a function of phyB activity (Table S3a), temperature (Table S3b), the interaction between phyB and temperature (Table S3c), and white light irradiance. The last term is used to incorporate the activity of the additional photo-sensory receptors (phyA, cry1 and cry2), which are not included in the term corresponding to phyB. To make the model more accurate under natural radiation conditions, where changes in spectral composition can alter the relative activation of different photo-sensory receptors, we modelled the specific contributions of phyA, cry1 and cry2 under controlled conditions. As in previous models, growth (G) is represented as a function of maximum growth (Go) divided by the sum of all the terms that reduce growth below the maximum (phyB, low temperature, interaction between phyB and low temperature, phyA, cry1 and cry2) plus 1 (Table S3); i.e., G=Go if there are no inhibitory terms.

To parameterise the term corresponding to phyA, we measured G both in WT and the *phyA* mutant 3-day-old light-grown seedlings exposed to a photoperiod of different mixtures of red plus far-red light, each one of them at a range of irradiances. Figure 1a shows the Go/G ratio for one of these red plus far-red light mixtures plotted against the sum of the rates of photoconversion from Pr to Pfr (k1) and from Pfr to Pr (k2). Go/G increases above 1 when there is inhibition (i.e G <Go). For a given spectral composition, k1+k2 increases with irradiance. The *phyA* mutant retained a significant degree of inhibition (equivalent to the phyB-mediated inhibition predicted by Legris et al. 2016) and the difference in slope between the WT and *phyA* provides an estimate of the inhibitory effect of phyA. Figure 1b shows the difference in slope between the WT and *phyA* observed in plots similar to that shown in Figure 1a but with different red / far-red ratios. The effect of spectral composition is captured by the k1/(k1+k2) ratio (we prefer the latter to the red / far-red ratio itself because the photoconversion rates are calculated with information provided by the whole spectrum (Mancinelli 1994)). The inhibitory effect is maximal for almost pure far-red light (the lowest k1/(k1+k2)) and steeply decreases with increasing proportions of red light to reach a plateau. This pattern is consistent with the well-known maximal effect of phyA under farred light (Quail *et al.* 1995), the contribution of phyA even under pure red light when the irradiance is high enough (Franklin *et al.* 2007) and the lack of effects of red / far-red ratio in the upper range of ratios (Sellaro *et al.* 2010). Then, the contribution of phyA was modelled by incorporating to the denominator a term where the activity of phyA depends on irradiance (*k1+k2*) in a manner that in turn depends on spectral composition, and more specifically red / far-red ratio (k1/(k1+k2)) (Table S3d).

To parameterise the terms corresponding to cry1 and cry2, we measured G both in WT and *cry1 cry2* mutant 3-day-old light-grown seedlings exposed to a photoperiod of different irradiances of blue light. Figure 1c shows the Go/G ratio plotted against blue-light irradiance (400-500 nm) and the difference in slope indicates the combined contribution of cry1 and cry2. To discriminate the role of each one of these receptors, we compared the double mutant to the *cry1* and *cry2* single mutants under a range of simulated sunlight and shade conditions and calculated the average ratio between G for each mutant and G for *cry1 cry2*. Figure 1d indicates strong genetic redundancy between cry1 and cry2 because the cry2 mutation on its own causes a reduction of G (reflected mathematically as “negative inhibition”) but significantly released the growth potential when combined with the *cry1* mutation. The contribution of cry1 and cry2 was modelled by incorporating a term where blue-light irradiance is multiplied by the slope of response when both cry1 and cry2 are present (Fig. 1d) multiplied by the specific contribution (Fig. 1e, Table S3e).

To test the performance of the model, seedlings of the WT and of the *phyA, phyB, phyA phyB, cry1, cry2* and *cry1 cry2*, were exposed to different simulated sunlight and shade conditions and different temperatures. Figure 1e shows the relationship between observed and predicted G, which did not significantly depart from the 1:1 line and showed adequate accuracy.

### 3.2 The impact of daily temperature fluctuations

While the model was able to capture the impact of temperature, the above experiments were conducted under conditions where this parameter remained constant through the photoperiod, which is not typical of natural settings. To investigate the impact of temperature fluctuations we first searched for natural patterns characterised by divergent daily trends and not very large differences in average temperature, in places with a photoperiod of 10 h (i.e. similar to the photoperiod used in our indoor experiments). We selected three locations in Argentina (Fig. 2a) and we exposed 3-day-old light-grown seedling of the WT and photo-sensory receptor mutants to a stable photoperiod in a growth chamber manually set to reproduce the three temperature patterns and a constant temperature control. We then measured G at different times of the photoperiod (Fig. 2b). The analysis of the residuals unaccounted by the growth model indicated that the accuracy of fit was improved by a factor reducing growth with increasing thermal amplitude in seedlings of the WT and of the *phyA* and *cry2* mutants (Table S4). This effect was not significant for the *phyB, cry1* and *cry1 cry2* mutants and was actually negative in the *phyA phyB* double mutant (Table S4). Thermal amplitude was calculated as the difference between maximum and minimum temperature with a negative signal if temperature decreases rather than decrease towards midday, as in the case of *Puerto Piramide* (Fig. 2a). Our interpretation of this effect is that in the WT (as well as *phyA* and *cry2*), during the photoperiod maximum hypocotyl growth rate occurs in the morning and then growth decreases gradually (Nozue *et al.* 2007; Sellaro *et al.* 2012). Therefore, if a given average temperature results from lower morning temperatures and higher values at midday, there is a coincidence between lower temperature and the time of maximal growth potential, which lowers average G. The *phyB, cry1, cry1 cry2* and *phyA phyB* mutants have higher rates not in the morning, but later in the photoperiod and therefore, do not become affected by temperature fluctuations in the same way. The model incorporating this additional term (Table S3f) accounted for the G values observed under fluctuating temperatures with reasonable accuracy (Fig. 1c).

### 3.3 The impact of daily light fluctuations

Following the same argument used for the analysis of fluctuating temperatures, we exposed 3-day-old light-grown seedling of the WT and photo-sensory receptor mutants to a photoperiod with irradiance manually adjusted hourly to obtain a peak at midday. We modified the red plus far-red light or the blue-light component leaving the other constant, or both components and we also included a control without temperature fluctuations (Fig. 3a). We then measured G during the first and second halves of the photoperiod (Fig. 3b). The model (which is based on average irradiance values) was able to predict G under these conditions with reasonable accuracy and no need for a correction associated to light fluctuations was apparent (Fig. 3c).

### 3.4 The impact of temperature or light fluctuations on PIF4 dynamics

The above experiments indicate that, compared to a stable condition with the same average temperature and irradiance, fluctuations in temperature peaking at midday tend to reduce growth, whilst fluctuations in irradiance do not have large effects. Since one of the most important players in the control of hypocotyl growth by light and temperature is PIF4 (Huq & Quail 2002; Niwa, Yamashino & Mizuno 2009; Franklin *et al.* 2011; Hornitschek *et al.* 2012), we investigated whether light and/or temperature fluctuations affect PIF4 dynamics. Figure 4 shows that compared to the stable conditions, increasing thermal amplitude significantly decreased PIF4 levels at midday. This observation offers a molecular correlate to the observed pattern of the physiological output.

**Figure 4.**
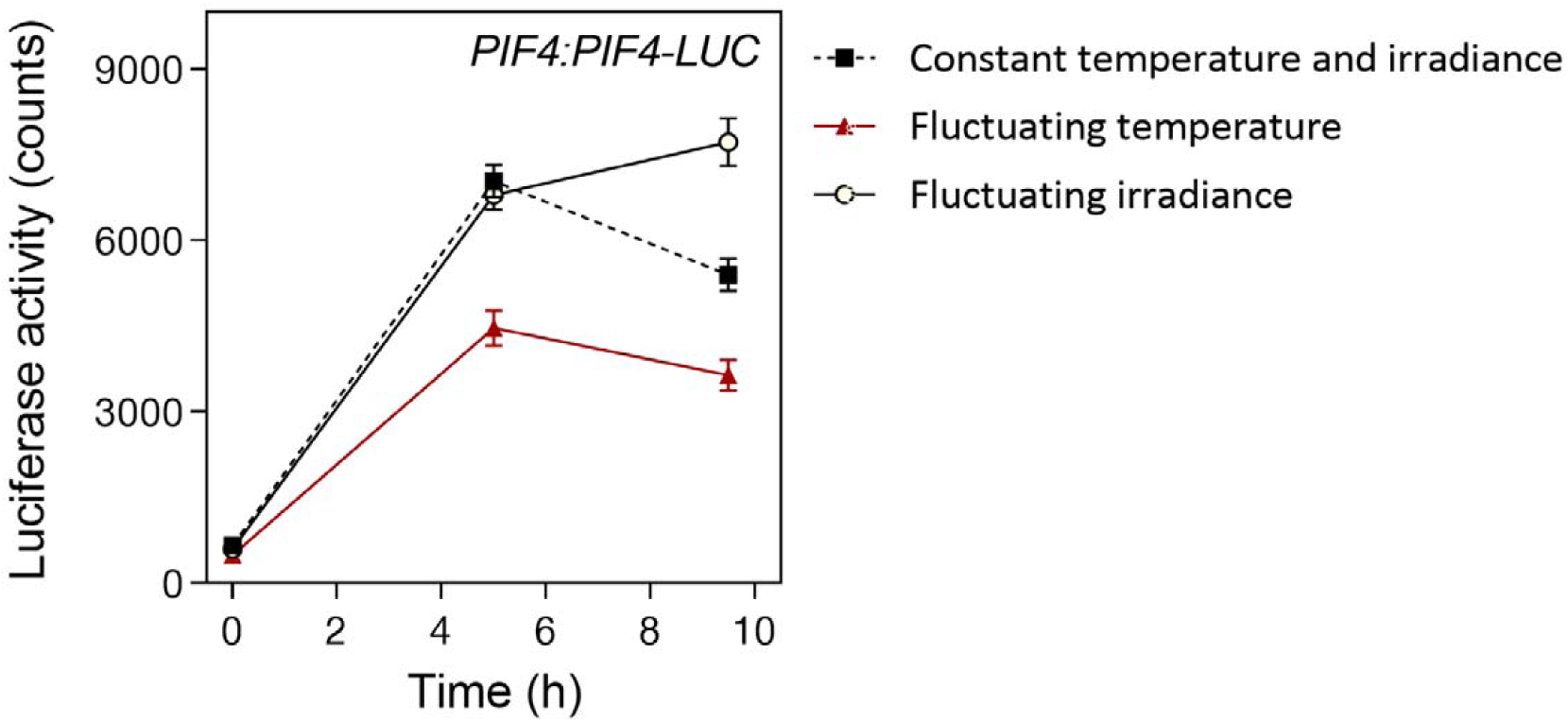
Thermal amplitude reduces the abundance of PIF4. Transgenic seedlings bearing the *PIF4:PIF4-LUC* transgene were exposed during the photoperiod of their fourth day to constant conditions of light (135 μmol. m^-2^. s^-1^) and temperature (20 °C), to fluctuating light and constant temperature, or to constant light and fluctuating temperature. Under fluctuating conditions, light and/or temperature peaked at midday and decreased (20 % per hour) towards the extremes of the photoperiod. Data are means and SE of three replicate plates.

### 3.5 Model performance in the field

To test the model developed under controlled conditions, we exposed 3-day-old light-grown seedling of the WT and photo-sensory receptor mutants to a photoperiod (10 h) of natural radiation either under full sunlight or under two levels of natural shade provided by grass canopies of different stature. The experiment was repeated on ten different dates between the beginning of winter and mid-summer to obtain a range of temperatures and irradiances. Light and temperature conditions inside the boxes where the plants were grown were monitored every hour. The daily average values were used as inputs to the model to estimate G (Table S3h). Figure 5 shows the relationship between observed and predicted data. The relationship did not departure significantly from the 1:1 ratio, showed no obvious bias due to genotype, light or temperature conditions and demonstrates that the model can predict G under field conditions with reasonable accuracy.

**Figure 5.**
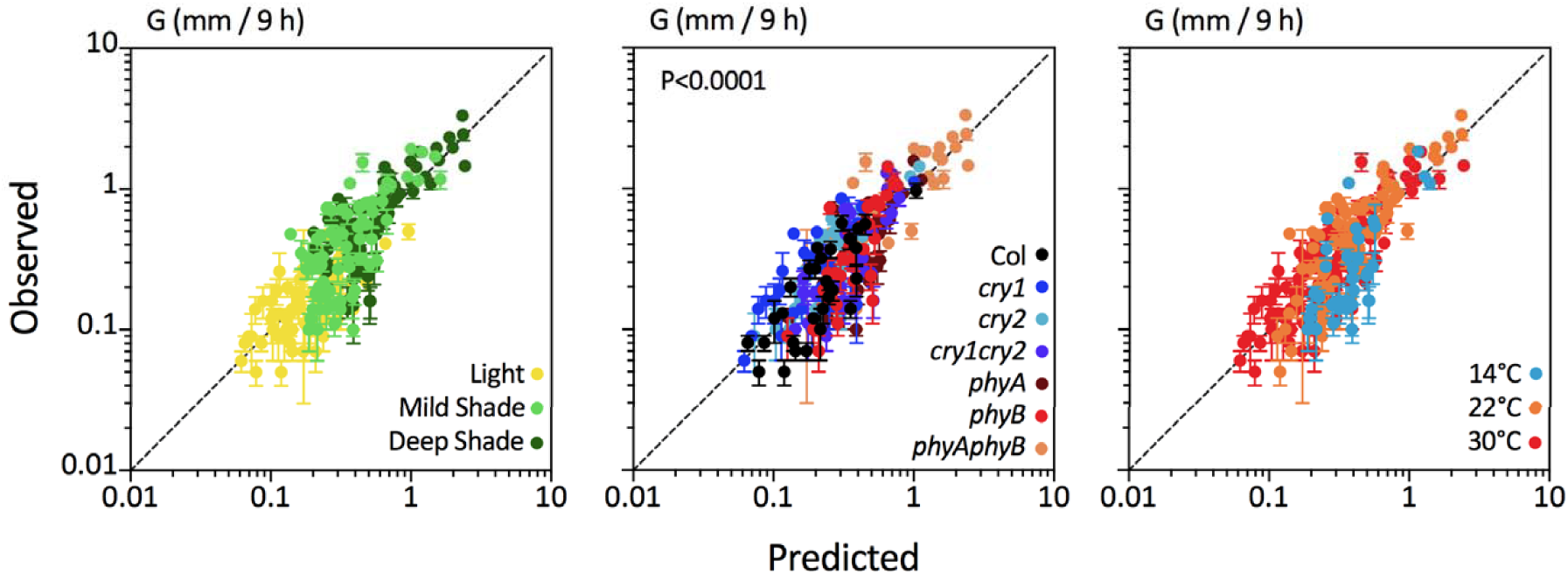
The model reasonably predicts hypocotyl growth in the field. Observed values of hypocotyl growth (G, Table S2) versus the values predicted by the model (Table S3h). Data are means and SE of four replicate boxes. The different light conditions, genotypes and temperatures are color-coded to show that the relationship between observed and predicted values is not biased for any of these factors (within the range tested here).

### 3.6 Impact of weather conditions on shade avoidance

To investigate the impact of weather conditions on shade avoidance, we first downloaded light and temperature data from 100 locations distributed over the Earth at the time of the year when the photoperiod is 10 h (Fig. 6a, Table S5). This photoperiod occurs only within certain latitudes, and very high latitudes are not included due to their extremely low temperatures (which are not suitable for Arabidopsis growth). Light and temperature data were used to estimate G under sunlight conditions, and corrected by the impact of the canopy to estimate G under mild and strong shade conditions. Figure 6b shows estimated G values under sunlight and shade conditions, plotted against the mean irradiance, mean temperature or thermal amplitude corresponding to each location (G values are the same in the three plots). The environmental values correspond to sunlight conditions (even though shade conditions were used to estimate G) because the aim is to explore the impact of weather. Of the three environmental variables, only mean temperature affected the magnitude of the shade avoidance response (the interaction between mean temperature and light/shade condition is significant at *P* < 10^−26^). The magnitude of the difference between G under sunlight compared with shade conditions increased under warmer temperatures.

**Figure 6.**
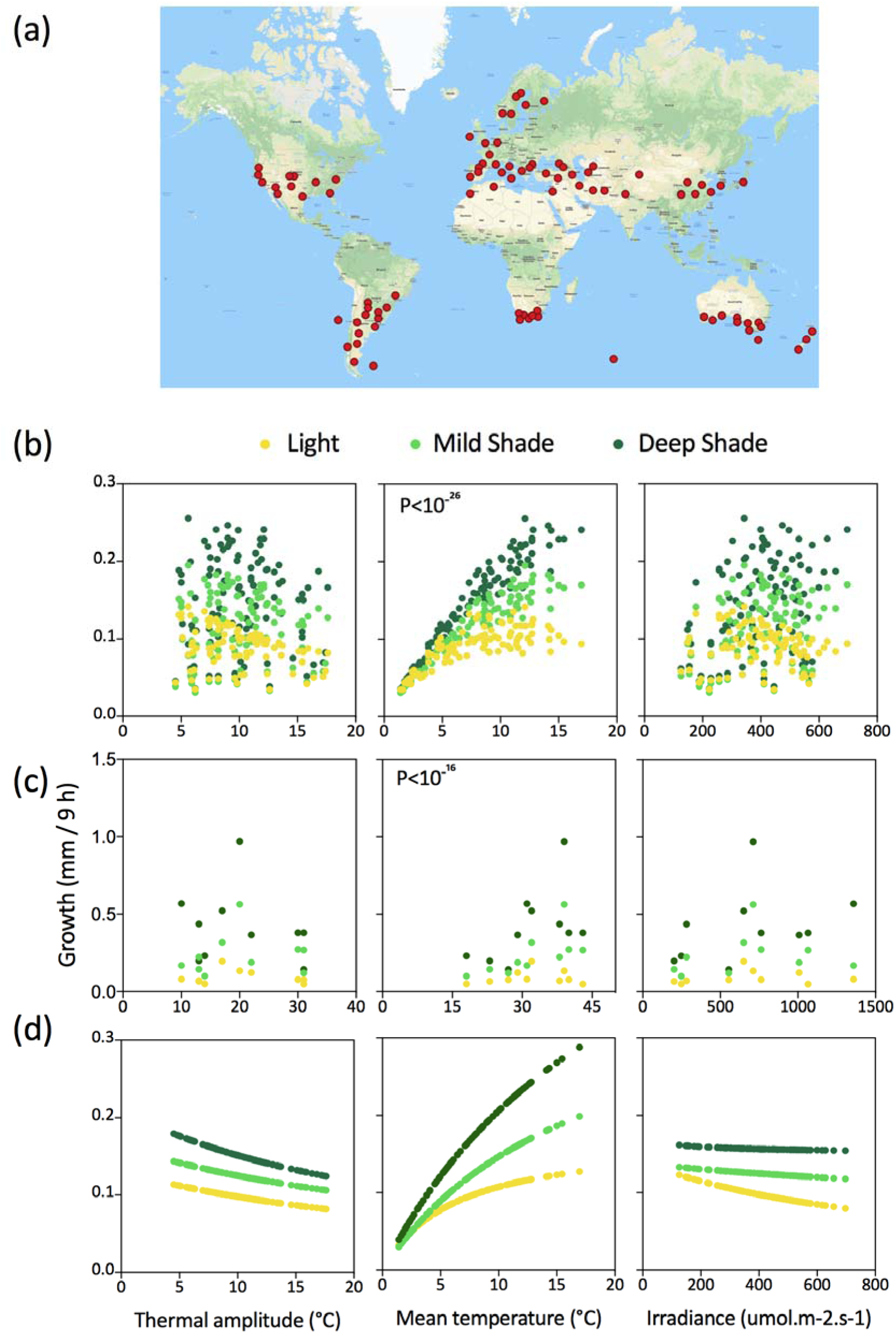
Warmer temperatures increase the shade-avoidance response. (a) One hundred locations for which weather data at a time of the year when photoperiod is 10 h was used as input for the model (Table S5). (b) Growth predicted by the model (Table S3h) for the different global locations against the thermal amplitude, average temperature or irradiance at that location. (c) Growth measured in field experiments (Table S2) against thermal amplitude, average temperature or irradiance at the date of the experiment. (c) Growth predicted by the model (Table S3h) for the thermal amplitude, average temperature or irradiance of a given location (the other two variables remain at the average value for all locations).

Figure 6c shows a similar plot but in this case, G corresponds to the values measured in our field experiments (Table S2, rather than model estimates) and their corresponding environmental data. Compared to the values observed in nature, the range is shifted towards higher temperatures because in our experiments the seedlings were exposed to natural radiation for 10 h even in the warm season, when natural photoperiod is above 10 h. Despite this, the results show the same pattern providing independent support to the conclusion based on the application of the model: Of the three environmental variables, only mean temperature affected the magnitude of the shade avoidance response (the interaction between mean temperature and light/shade condition is significant at *P* < 10^−16^).

Finally, in Figure 6d, G was again estimated by using the model and the variables corresponding to the 100 world locations but in this case only the variable plotted in abscissas was varied, whereas for the other two, the average for the 100 locations was used as input. This procedure eliminates the natural correlations among temperature, irradiance and thermal amplitude. The results of this exercise show that irradiance and thermal amplitude theoretically do have impact on shade avoidance; which becomes more intense under better irradiated conditions with weaker thermal amplitude. The comparison of Figs 6a and 6c indicates that the effect of irradiance and thermal amplitude are not obvious in natural settings because they become diluted and/or counteracted by variations in the other variables.

### 3.7 Predicted impact of climate change on the magnitude of shade-avoidance responses

By using current weather data in combination with a model that estimates climate trends, we obtained predicted whether conditions for 2080 at a global scale. These data were used as input for the growth model (Table S3h) to estimate current and future growth under sunlight, and two different degrees of shade. In the plot of future vs current growth, most of the points are above the 1:1 line (Fig. 7). This pattern is consistent with the fact that the elevation of average temperature with global warming increases hypocotyl growth. The extent of change (future minus current estimated growth) is significantly higher for the seedlings grown under deep shade, compared to mild shade or sunlight (Fig. 7, inset). This indicates that shade-avoidance responses are predicted to increase in a scenario of global warming. It must be emphasised, however, that this exercise is valid only to investigate the impact of future combinations of temperature, irradiance and thermal amplitude, whilst a prediction of the actual growth rates would require the incorporation of other variables to the model (e.g. water availability).

**Figure 7.**
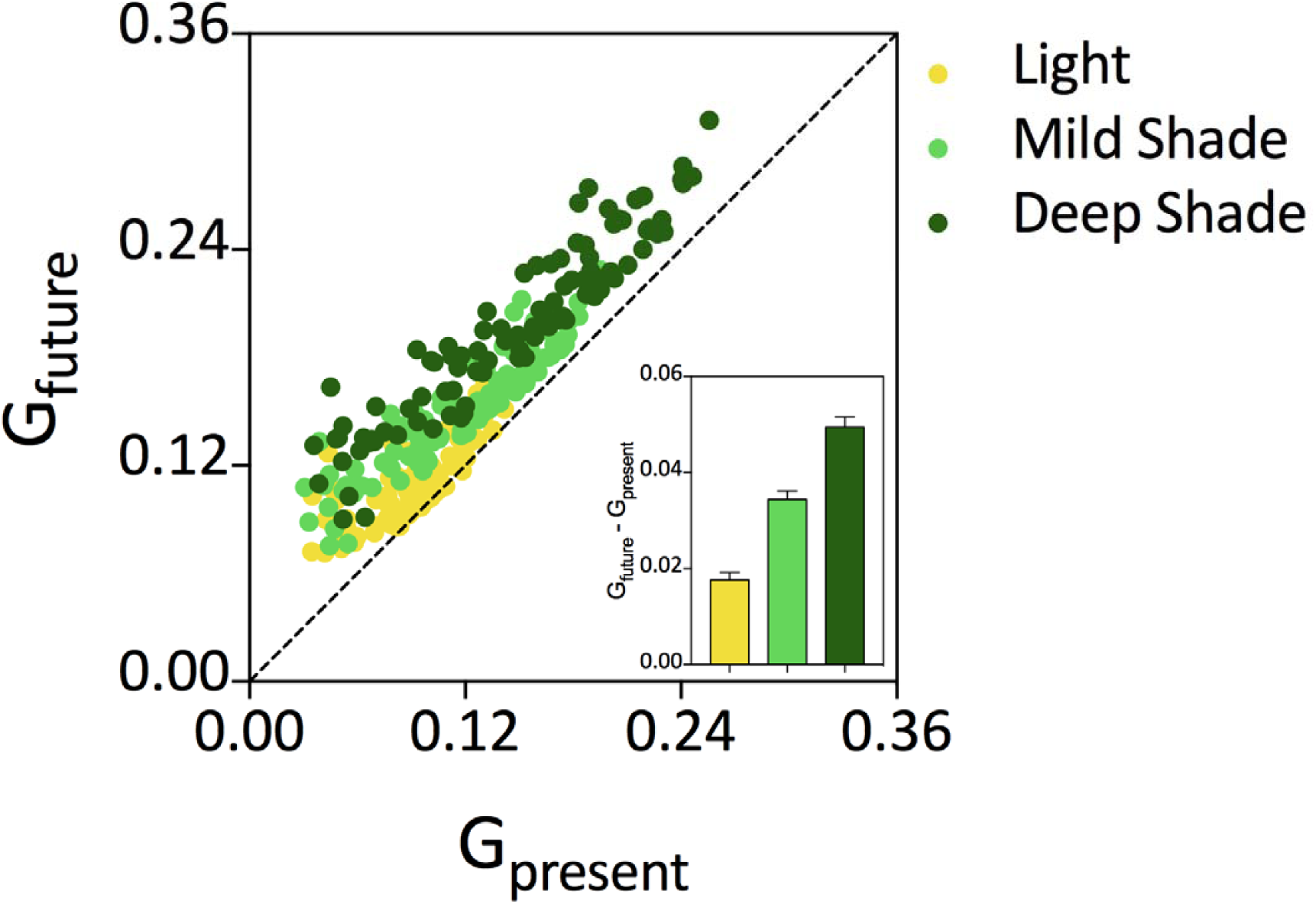
Predicted enhancement of the shade-avoidance response in a scenario of global change. Growth rate was calculated by using the model (Table S3h) in combination either with current or estimated (future) weather data. The inset shows the average difference between future and current growth averaged for all locations in plants grown under sunlight, mild shade or deep shade.

## 4 DISCUSSION

Our understanding of the mechanisms involved in sensing the degree of shade or temperature as well as of the nature of the downstream signalling steps has progressed substantially in recent years (see Introduction for references). In this context, it is fair to ask how accurately this knowledge, obtained largely under controlled conditions, accounts for the patterns of plant growth in the field. Here we present a model that reasonably predicts hypocotyl growth in *A. thaliana* seedlings during the photoperiod as a function of field conditions of light and temperature (Fig. 5). The structure of the model resembles that used for previous hypocotyl growth models (Rausenberger *et al.* 2010; Sellaro *et al.* 2010; Legris *et al.* 2016) as it is based on the maximum growth values divided by terms that represent the conditions that reduce growth below the maximum (Table S3h). However, it differs from previous models because it predicts growth rate within a specific time window rather than final hypocotyl length (Rausenberger *et al.* 2010; Sellaro *et al.* 2010), it incorporates light and temperature inputs as opposed to models based on light conditions only (Rausenberger *et al.* 2010; Sellaro *et al.* 2010), and/or it dissects phyB, phyA and cry activity rather than phyB alone or a general term for various photo-sensory receptors (Rausenberger *et al.* 2010; Legris *et al.* 2016). In addition, the model incorporates several interactions between light and temperature conditions that fine-tune the predictions.

When compared to the conditions normally used in growth chambers, one of the distinctive features of the natural environment is the fluctuation of light and temperature, which often peak close to midday. Our analysis did not reveal a significant impact of the light fluctuations (i.e., the daily light average was a sufficiently accurate input, Fig. 3). However, we did observe that hypocotyl growth rate decreased with thermal amplitude (Fig. 2). A reasonable interpretation of this observation could be based on the fact that faster hypocotyl growth rates occur in the morning (Nozue *et al.* 2007; Sellaro *et al.* 2012). Larger thermal amplitudes would imply lower morning temperatures; therefore, growth would be limited by low temperatures at the time of maximum potential. Thermal amplitude decreased the abundance of PIF4 (Fig. 4), which is a key driver of hypocotyl growth (Huq & Quail 2002; Niwa *et al.* 2009; Franklin *et al.* 2011; Hornitschek *et al.* 2012).

We applied the model to investigate the impact of weather conditions (incoming irradiance, average temperature, thermal amplitude) on the hypocotyl growth response to shade. We used weather conditions typical of different locations of the world at the time of the year when the photoperiod is *ca*. 10 h (Fig. 6a). The latter is the photoperiod under which the model was developed under controlled conditions and tested in the field. The model indicates a significant positive effect of average temperature on the extent of response to shade (Fig. 6b). Meanwhile, neither incoming irradiance nor thermal amplitude affected the magnitude of shade avoidance. Although more restricted in terms of spread of weather conditions, the actual field data supports the same conclusion (Fig. 6c). Therefore, for a given genotype, shade-avoidance responses tend to be more intense at warmer locations of the Earth. A different issue that remains to be elucidated is whether there is adaptation and the genotypes typical of warmer places are more or less responsive to shade than those from cooler areas. It must also be noted that the model does not incorporate other environmental conditions (water, nutrients), which can also limit growth. A stronger shade avoidance under warm conditions would reduce the double jeopardy caused by the low light input for photosynthesis under shade plus the high consumption of carbohydrates in respiration under high temperatures (Casal & Balasubramanian 2019).

There are previous reports showing that plants may show stronger responses to neighbour signals at warmer than at cooler temperatures when light and temperature conditions are manipulated independently (Wall & Johnson 1982; Mazzella *et al.* 2000; Weinig 2000; Halliday & Whitelam 2003; Patel *et al.* 2013). One could therefore ask whether the model approach was actually necessary in the first place. The observations reported here provide a conclusive answer to this question. The model approach was necessary because under natural conditions light and temperature features of the environment do not fluctuate in a fully independent manner (Legris, Nieto, Sellaro, Prat & Casal 2017). As a result of the latter, the impact of one variable can be compensated by the effects of another variable correlated with the former. In fact, if we change one variable at the time (i.e. leaving the others at a constant value) shade avoidance responses should be affected not only by average temperature, but also by irradiance and thermal amplitude (Fig. 6d). The impacts of irradiance and thermal amplitude only disappear when we use the actual records of these variables for different locations, which imply the natural correlations among environmental variables.

Our model predicts that as a result of global warming shade-avoidance responses would tend to increase in the future (Fig. 7). Of course, other aspects of global climate change (e.g. water availability) are not considered by the current growth model and could distort this trend. Bearing in mind this limitation and taking into account that shade-avoidance responses have both negative (Robson, McCormac, Irvine & Smith 1996) and positive effects (López Pereira, Sadras, Batista, Casal & Hall 2017) on crop productivity, the sensitivity to neighbour cues might need to be modified to optimise crop plant architecture in the context of global climate change.

## ACKNOWLEDGEMENTS

We thank Salomé Prat (*Centro Nacional de Biotecnología*, Madrid, Spain) for her kind provision of Arabidopsis seeds carrying the *PIF4:PIF4-LUC* transgene. This work was supported by a joint proposal funded by *Deutscher Akademischer Austauschdienst* (DAAD) and the Secretariat of Science of Argentina (grant DA/16/04 to M.D.Z and J.J.C.), the Deutsche Forschungsgemeinschaft (grant ZU259/2-1 to M.D.Z.) and under Germany’s Excellence Strategy CEPLAS EXC2048/1 ID 390686111 (to M.D.Z.), the University of Buenos Aires (Grant 20020100100437 to J. J. C.), and Agencia Nacional *de Promoción Científica y Tecnológica* (Grant PICT-2016-1459 to J. J. C.).

## SUPPORTING INFORMATION

**Figure S1.** Spectral photon distribution of the light under controlled conditions.

**Table S1**. Hypocotyl growth data under controlled conditions

**Table S2.** Hypocotyl growth data under field conditions.

**Table S3.** Parameters of the growth model

**Table S4.** Statistical analysis of thermal fluctuation terms.

**Table S5.** Weather data at different global locations.

