## Supplemental Table 3 for "Shade-avoidance responses become more aggressive in warm environments"

| **Description** | **Term** | **Values** |
| --- | --- | --- |
| a. Effect of phyB | *k. D2* | *D2* is calculated from the light scans.  *k*=3.6 (Legris et al., 2016) |
| b. Effect of low temperatures | *a* | *a=50* (Legris et al., 2016)  *a=* 21.3 in the *phyA phyB* mutant  T=average temperature |
| c. Interaction between phyB and low temperatures | *c* | *c*=63.24 (Legris et al., 2016) |
| d. Effect of phyA | *d . (k_1_+k_2_) . PHYA* | *k1+k2* is calculated from the light scans. *PHYA=1* if phyA is present or *=1* in the *phyA* mutant.  $d=721 \times e^{-32 \times k1/(k1+k2)}-4.059$ |
| e. Effects of cry1 and cry2 | *e* . *B_400-500 nm_* . *CRY* | *e*= *0.048* (slope of response to blue light in the WT).  *e*= *0.0* if temperature is >28°C.  *B_400-500 nm_* is the blue-light irradiance (400-500 nm).  e=0.22 in the *phyA phyB* mutant when T <22°C).  =0.04 in the *phyA phyB* mutant when T >22°C)  *CRY*= 1 if both cryptochromes are present, 0.76 if only cry2 is present, 1.3 if only cry1 is present and 0 if both are absent. |
| f. Effect of thermal amplitude | *f. AT* | *f*=0.7 for WT, *cry2* and *phyA*.  *f*=0.21 for *cry1* and *phyB.*  *f*=0 for *cry1 cry2* and *phyA phyB*.  AT= thermal amplitude (maximum minus minimum) |
| g. Maximum growth rate | *G_0_* | *G_0_*=2.30  *G_0_*= 3.07 in the *phyA phyB* mutant  We observed a higher growth potential than that reported by Legris et al. (2016), likely attributable to the fact that here the plants were grown at 100 µmol.m^-2^.s^-1^ before the treatments, compared to the 50 µmol.m^-2^.s^-1^ used by Legris et al (2016)*.* |

Table S3. Parameters of the growth model

h. Structure of the growth model:

$$\text{G}\text{ }\text{= }\frac{G_{0}}{\text{1 + }\text{a}\text{ . }\left( \frac{\text{1}}{\text{T}}\text{-}\frac{\text{1}}{\text{30°C}} \right)\text{+ }\text{c}\text{ . D2 . }\left( \frac{\text{1}}{\text{T}}\text{-}\frac{\text{1}}{\text{30°C}} \right)\text{+ }\text{d}\text{ . }\left( \text{ }\text{k1+k2} \right)\text{. PHYA + }\text{e}\text{ . }B_{400-500 nm}\text{ . CRY + }\text{f}\text{ . AT}}$$
