## Supplemental Table 4 for "Shade-avoidance responses become more aggressive in warm environments"

|  | *Coefficients* | *Probability* |
| --- | --- | --- |
| Intercept | 0,00 | #N/A |
| AT . cry1 | 0,21 | 0,02 |
| AT . (WT/cry2/phyA) | 0,70 | 0,00 |
| AT . phyB | 0,21 | 0,01 |
| AT . cry1 cry2 | -0,08 | 0,29 |
| AT . phyA phyB | -0,04 | 0,14 |

Table S4. Statistical analysis of thermal fluctuations terms.
