## Supplementary figures and images for "Shade-avoidance responses become more aggressive in warm environments"

### Supplemental Figure 1

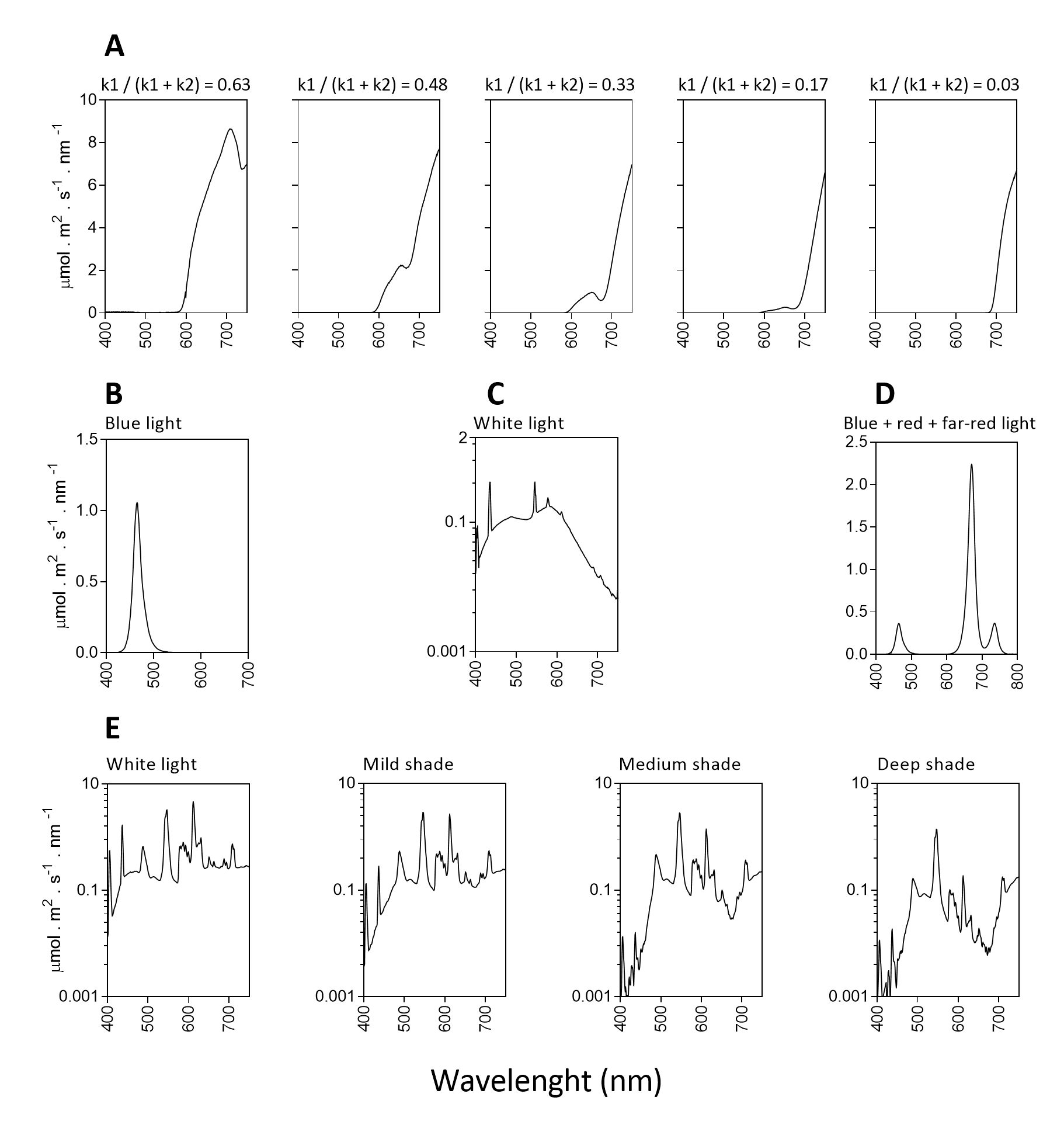
